# The Spread of Interferon-γ in Melanomas is Highly Spatially Confined, Driving Non-Genetic Variability in Tumor Cells

**DOI:** 10.1101/2023.01.26.525713

**Authors:** Edoardo Centofanti, Chad Wang, Sandhya Iyer, Oleg Krichevsky, Alon Oyler-Yaniv, Jennifer Oyler-Yaniv

**Affiliations:** The Department of Systems Biology at Harvard Medical School, Boston, Ma 02115; The Systems, Synthetic, and Quantitative Biology Graduate Program at Harvard Medical School, Boston, Ma 02115; The Department of Physics at Ben Gurion University of the Negev, Beer-Sheva Israel

## Abstract

Interferon-γ (IFNγ) is a critical anti-tumor cytokine that has varied effects on different cell types. The global effect of IFNγ in the tumor depends on which cells it acts upon and the spatial extent of its spread. Reported measurements of IFNγ spread vary dramatically in different contexts, ranging from nearest-neighbor signaling to perfusion throughout the entire tumor. Here, we apply theoretical considerations to experiments both *in vitro* and *in vivo* to study the spread of IFNγ in melanomas. We observe spatially confined niches of IFNγ signaling in 3-D mouse melanoma cultures and human tumors that generate cellular heterogeneity in gene expression and alter the susceptibility of affected cells to T cell killing. Widespread IFNγ signaling only occurs when niches overlap due to high local densities of IFNγ-producing T cells. We measured length scales of ∼30-40μm for IFNγ spread in B16 mouse melanoma cultures and human primary cutaneous melanoma. Our results are consistent with IFNγ spread being governed by a simple diffusion-consumption model, and offer insight into how the spatial organization of T cells contributes to intra-tumor heterogeneity in inflammatory signaling, gene expression, and immune-mediated clearance. Solid tumors are often viewed as collections of diverse cellular “neighborhoods”: our work provides a general explanation for such non-genetic cellular variability due to confinement in the spread of immune mediators.

## Introduction

Interferon-γ (IFNγ) is a broadly acting cytokine with important anti-tumor effects that is produced primarily by activated CD4+ and CD8+ T cells ^1^. IFNγ mediates anti-tumor immunity during surveillance for occult cancers ^2^ and regression of established tumors ^3–6^. Moreover, the effectiveness of checkpoint blockade therapies relies on an intact IFNγ signaling axis ^7–9^.

The anti-tumor effects of IFNγ are varied and exerted on resident immune cells, stromal cells, and tumor cells. IFNγ acts directly on tumor cells ^10–13^ and stromal cells ^5,6,14^ to cause cytotoxicity, mitotic arrest, and senescence. These effects play an important role in tumor clearance because contact-dependent elimination of tumor cells by cytotoxic T cells is relatively inefficient ^12^. IFNγ acts on immune cells to polarize them towards a more inflammatory state ^15^ and lowers the activation barrier for T cells to respond to antigen ^16^. IFNγ also upregulates tumor cell antigen presentation, thereby priming the cells for improved recognition by T cells ^14,17^. The specific effects of IFNγ depend on the identity and number of cells it acts upon, and by extension, on its spatial perfusion through the tumor. However, it is unclear how far IFNγ spreads away from its source of production, and what factors govern this spread. Different studies have yielded contradicting results.

Some studies have concluded that IFNγ is highly spatially confined because it is secreted directionally into the immune synapse, thereby signaling only to the T cells’ cognate antigen presenting cell ^18–22^. Other work reports that IFNγ leaks beyond the immune synapse, but only signals to cells in the local vicinity ^23,24^. In such contexts, steep gradients of IFNγ signaling are expected: cells in immediate proximity to the producing T cell will be subject to high IFNγ concentrations, while the majority of the tissue will not be exposed to IFNγ at all. Conversely, IFNγ has been reported to spread widely throughout an infected tissue ^25,26^ or tumor ^27,28^, acting on bystander cells that are on the order of hundreds of microns or more away from the assumed source of production. These bystander effects of IFNγ have been proposed to exert long-term changes to tumor gene expression ^27^ and serve as a fail-safe mechanism in situations where antigen-loss variants become resistant to T cell killing ^28^. These diverse findings have yet to be reconciled, motivating us to investigate the fundamental principles that control cytokine spread ^29,30^.

We and others previously established that the spatial spread of many diffusible growth factors and cytokines away from their source is governed by simple diffusion and consumption dynamics ^31–34^. Accordingly, the length scale of cytokine spread, defined as the average distance traveled by a cytokine molecule, is determined entirely by three parameters: *(1)* the molecular diffusion rate, *(2)* the density of cells capable of consuming the cytokine, and *(3)* the quantity of receptors on cytokine consuming cells. We used this simple quantitative framework to measure the spread of Interleukin-2 in lymphoid organs ^31^, but similar conceptual models have since been used to explain the spread of TNF-α ^35^, PGDF and CSF-1 ^36^, chemokines ^37,38^, and type I Interferon ^39^. High, ubiquitous cytokine receptor expression limits the spatial spread of signaling and generates niches of altered gene expression and localized regulation of immune responses, whereas low levels of receptor expression leads to more uniform signaling across a tissue ^36,40–43^. This conceptual and quantitative framework is a broadly useful and flexible approach to study the spatial regulation of diffusible cytokines in densely-packed, three-dimensional (3-D) tissues.

We applied this diffusion-consumption framework to study the spatial spread of IFNγ *in silico*, through mouse B16-F10 (B16) melanoma cells *in vitro*, and in melanoma specimens taken from human patients. We discovered that the spread of IFNγ in these situations is, in fact, spatially confined to small niches of cells surrounding an IFNγ-producing T cell. This spatial confinement leads to significant non-genetic cellular variability in downstream signaling through STAT1 and IRF1, antigen presentation, and ultimately impacts T cell mediated killing. Our study demonstrates that the frequency and spatial distribution of tumor infiltrating T lymphocytes (TILs) are critical factors determining the degree of perfusion of melanomas by IFNγ. We propose that in tumors expressing IFNγ receptors, which appear common in mouse and human melanomas, widespread IFNγ signaling is only achieved when the density of TILs is high enough that localized cytokine fields overlap. In other types of cancers, or diseased tissues, IFNγ receptor expression may be lower, leading to a wider spatial distribution of IFNγ. Our work reinforces the notion of tumors as heterogeneous collections of cellular “neighborhoods’’ by providing a general mechanism that spatially restricts the spread of immune mediators to confined regions.

## Results

### TIL density varies dramatically both within individual tumors and between patients

IFNγ originates from activated lymphocytes, so we first examined the spatial distribution of tumor infiltrating lymphocytes (TILs). To do so, we studied published FFPE tumor biopsies taken from several anatomical sites of a patient with state II nodular melanoma ^44^. The tumor lesions were multiplex antibody stained using cyclic immunofluorescence microscopy (CyCIF) ^45,46^. Tumor cells were identified by their expression of S100A - a standard melanoma marker ^47^ - and T cells were identified by their expression of CD3, CD4 or CD8, and absence of FoxP3 (Fig S1A). We further defined TILs as T cells that are within 25μm of at least 10 tumor cells (Fig S1B). This eliminates T cells that exist outside the tumor boundary, which are not, by definition, infiltrating the tumor. We then re-constructed the images such that S100A+ cells are represented as gray points and TILs are represented as red points (Fig 1A). These images demonstrate vastly different overall tumor infiltration by T cells in different sites, with the *average* TIL density ranging from ∼1 in 300 to ∼1 in 30 tumor cells. Moreover, even in a tumor sample with a high *average* TIL density (1 TIL in 87 tumor cells), the *local* TIL density can be extremely variable (Fig 1B-C). The local relative density of TILs - represented by color intensity in Fig 1C - was computed using a gaussian kernel density estimator with a 100μm bandwidth. We observed regions of this tumor with very high local TIL density of ∼1 in 10 tumor cells, while other areas were only sparsely populated by TILs or devoid of them altogether (Fig 1B-E). In this tumor - representing a sample with relatively high *average* TIL density, nearly 25% of the tumor has a *local* TIL density equal to or less than one TIL in one thousand tumor cells (Fig 1D).

To translate the ratio of TILs to tumor cells to a distance between TILs, we computed the theoretical mean inter-particle distance between TILs in the context of a densely-packed 3-D space (Fig 1F). This calculation incorporates simplifying assumptions that, locally, TILs are distributed randomly, and that the typical diameter of a cell is 10μm. In regions where TIL density is one in ten thousand cells, TILs are spaced on average ∼13 cell diameters (130μm) apart. In regions where TIL density is one in one hundred cells, however, TILs are spaced on average ∼3 cell diameters (30μm) apart, indicating that most tumor cells have at least one nearest neighbor that is a TIL.

**Figure 1:**
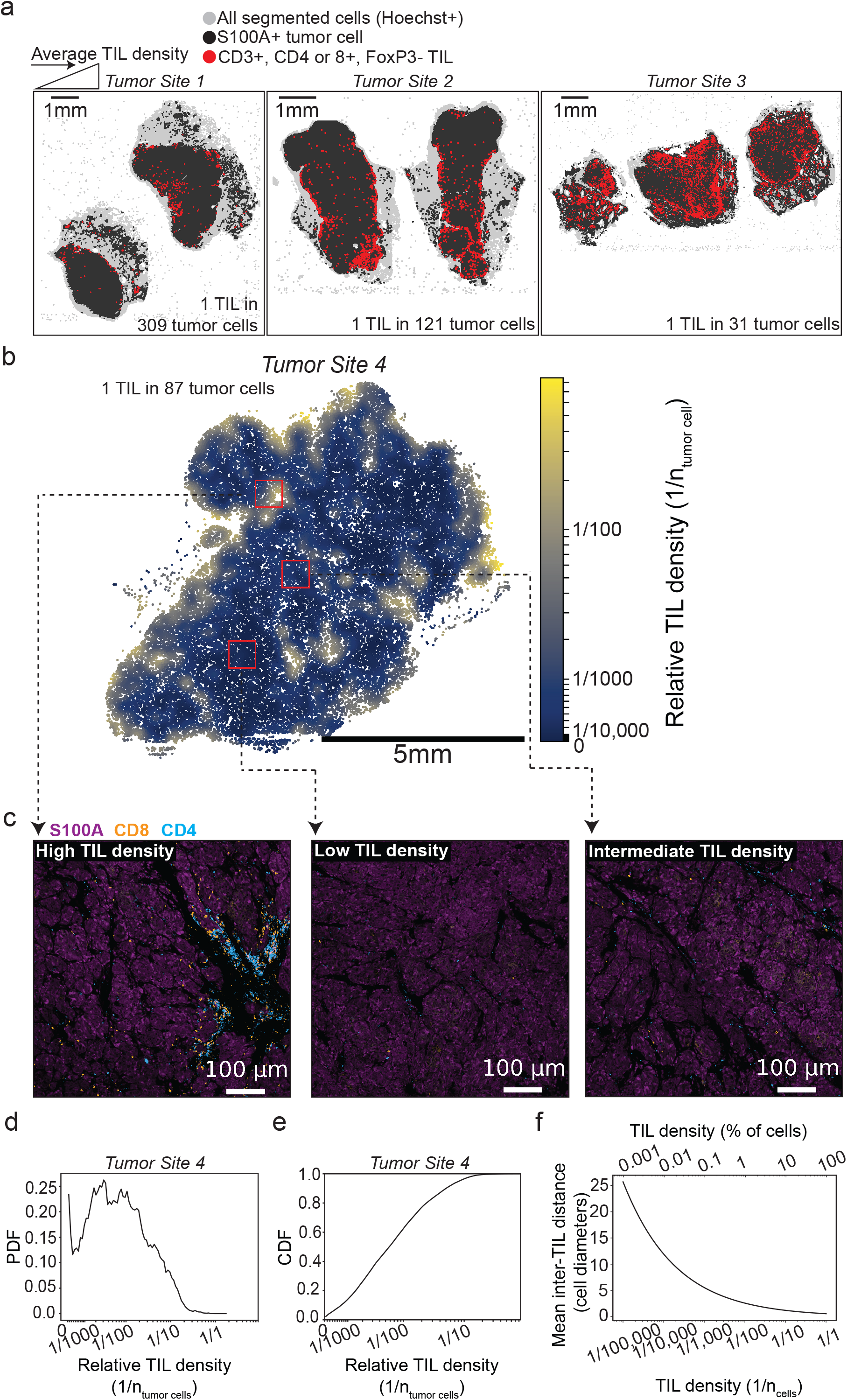
TIL density varies dramatically within individual tumors and across different tumors. **(a)** Image reconstructions from melanoma lesions stained and imaged using CyCIF and obtained from three different sites of one patient. Single cells were first segmented using Hoechst (nuclear) staining, and tumor cells were assigned by thresholding on the level of S100A staining. T cells were identified by thresholding on levels of CD3, CD4 or CD8, and absence of FoxP3. Infiltrated T cells (TILs) were further selected by removing all T cells not within at least 25μm from at least 10 S100A+ tumor cells. Images were then converted to point clouds where tumor cells are represented by gray points and TILs are red. **(b)** Local relative TIL density in a melanoma lesion from an additional tumor site. Local TIL density was computed by scanning over the image with a 100μm bandwidth gaussian kernel density estimator. **(c)** Images demonstrating variable TIL density from each of the red highlighted regions in (b). **(d)** Probability distribution function of local TIL density depicted in (b). **(e)** Cumulative distribution function of local TIL density depicted in (b). **(f)** Theoretical inter-particle distance between TILs in a dense-packed 3-D environment as a function of TIL density. We assume randomly scattered TILs and that all cells are 10μm in diameter.

These analyses demonstrate dramatic spatial heterogeneity in TIL-tumor cell interactions within individual tumors and across different tumor sites. For IFNγ to perfuse an entire tumor given this heterogeneity, it would need to diffuse over hundreds of microns. If IFNγ instead spreads only a short distance, such heterogeneity in TIL spacing would generate significant regional variability in tumor IFNγ signaling and gene expression. To distinguish between these two scenarios, we next measured IFNγ signaling in melanoma cells cultured in a dense, 3-D environment.

### Dense, 3-D culture conditions generate cell-to-cell heterogeneity in the response to IFNγ

One of the key challenges in studying cytokine spread *in vitro* is that it is hard to mimic the high cell density and 3-D geometry that exists in tissues. Conventional tissue culture plates culture cells at low density and lack 3-D structure, and gradient formation is limited by convective flows that mix cytokines in the media. We previously established the clusterwell plate, a custom-fabricated well plate that better approximates *in vivo* conditions, as a method for generating *in vivo*-like cytokine gradients *in vitro* (Fig 2A) ^31^.

**Figure 2:**
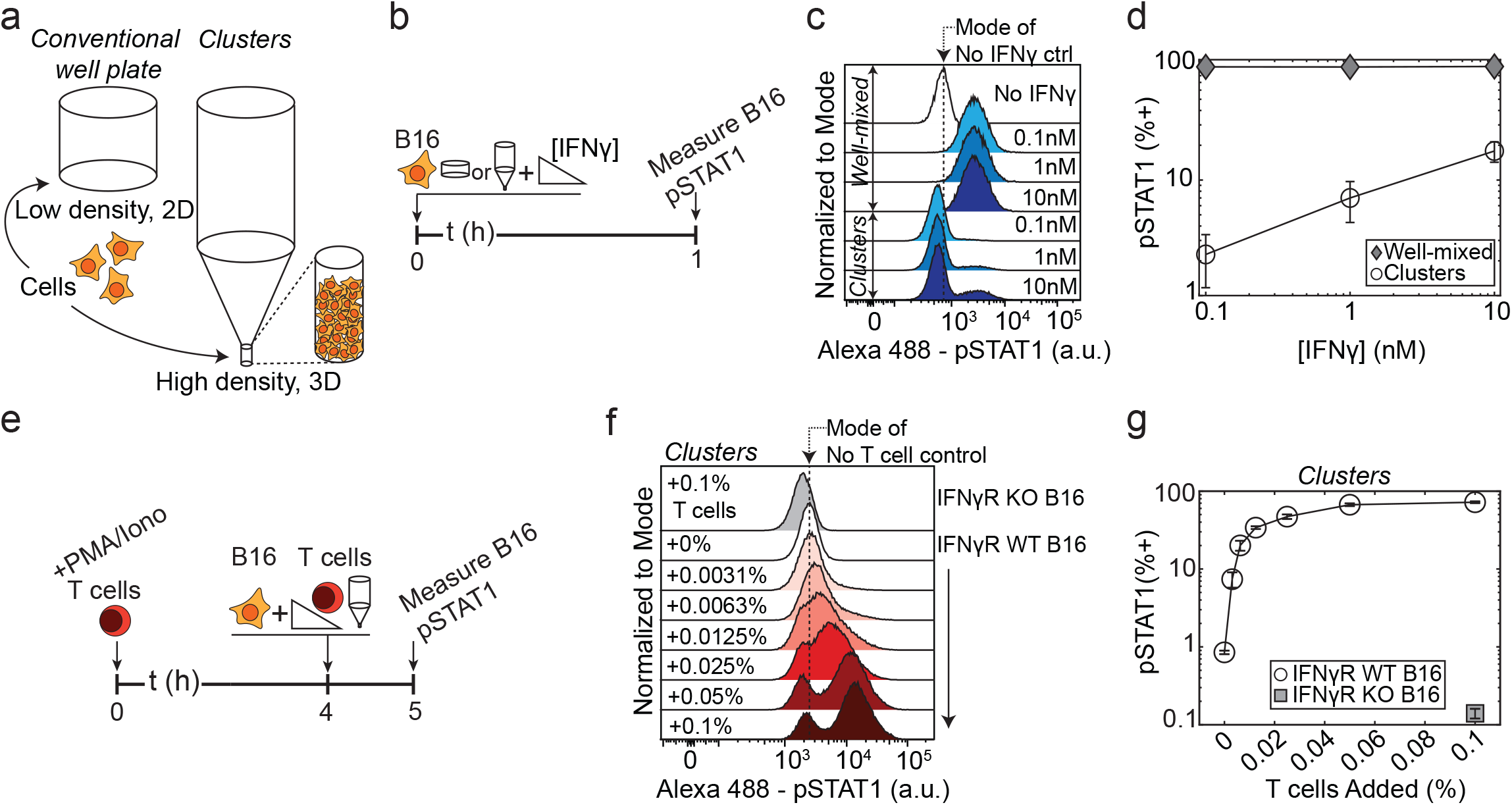
Dense, 3-D culture conditions generate cell-to-cell heterogeneity in the response to IFNγ. **(a)** Diagram of the basic geometry of conventional well plates and our clusterwell plates. Clusterwells culture cells in 3-D and at high density compared to well plates. **(b)** Experiment diagram for results shown in (c-d). B16 cells were cultured in either conventional 96 well plates or in clusters and stimulated with either 0.1, 1, or 10nM of IFNγ for 60m, before staining for pSTAT1 and performing flow cytometry. **(c)** Histograms of pSTAT1 levels in IFNγ-stimulated B16 cells. **(d)** Percentages of pSTAT1+ cells from (c). **(e)** Experimental diagram for results shown in (f-g). T cells were stimulated with PMA and Ionomycin for 4h before being co-cultured at varying densities with B16 cells in clusterwells for 1h. Cells were then stained for pSTAT1 and measured by flow cytometry. **(f)** Histograms of pSTAT1 levels in B16 cells co-cultured with activated T cells. **(g)** Percentages of pSTAT1+ cells from (f).

We used our clusterwells to evaluate how a densely-packed 3-D environment affects IFNγ signaling in a population of B16 mouse melanoma cells. To determine whether steep IFNγ gradients form in dense environments, we loaded the cells into the clusterwells and added exogenous IFNγ on top. We compared the signaling response in the clusterwell to cells that were cultured in conventional 96 well plates (denoted “well-mixed” onward) (Fig 2B). IFNγ signaling was evaluated by staining for phosphorylated STAT1 (pSTAT1), the transcription factor immediately downstream of the IFNγ receptor ^1^, and performing flow cytometry.

In well-mixed conditions, B16 cells exhibit a graded, dose-dependent, unimodal response to IFNγ (Fig S2A-B, 2C-D). In contrast, B16 cells cultured in clusterwells show a bimodal response to IFNγ where the proportion of pSTAT1+ cells scales dose-dependently with IFNγ (Fig 2C-D) ^31^. Thus, even at high concentrations, IFNγ does not fully penetrate the cluster of cells, leading to spatial heterogeneity in cytokine exposure. These results were identical in cultures stimulated for longer periods of time (90 min), indicating that IFNγ signaling had reached steady state (Fig S2C-D). We previously showed that the extent of cytokine penetration into cell clusters depends on cytokine consumption, rather than molecular degradation ^31^.

To determine whether IFNγ signaling remains bimodal when the cytokine is produced by cells intermixed with the target cells, we co-cultured B16 cells with different numbers of activated, IFNγ-producing T cells in our clusterwells (Fig 2E). We activated CD8+ OT-1 T cells for four hours, causing the vast majority of them to produce IFNγ (Fig S2E-F). OT-1 T cells bear an Ovalbumin (OVA)-specific transgenic T cell receptor (TCR) that is non-reactive towards B16 cells in the absence of OVA antigen, and can thus be used purely as a source of IFNγ. Again, we observed a bimodal pSTAT1 response where the proportion of pSTAT1+ cells scales dose-dependently with the density of activated T cells (Fig 2F-G). This STAT1 signaling is IFNγ-dependent; IFNγ receptor knockout (IFNγR KO) B16 cells did not phosphorylate STAT1 despite co-culture with the highest density of activated T cells. These data indicate that spatial effects in our clusterwell cultures lead to differential exposure of target cells to IFNγ, which in turn leads to cellular heterogeneity in phosphorylation of STAT1.

### Imaging reveals niches of IFNγ signaling surrounding activated T cells, and enables direct measurement of the signaling length scale

To visualize and measure the formation of IFNγ gradients directly, we imaged the pSTAT1 response of OVA-pulsed B16 cells co-cultured with OT-1 T cells using our established PlaneView imaging approach ^31,48^. Briefly, we mixed OVA-pulsed B16 cells with CFSE-labeled OT-1 T cells and deposited them in a monolayer on a fibronectin coated glass slide (Fig 3A). Then, 10 layers of B16 cells were added on top, forming a 3-D array with T cells restricted to the bottom layer. The co-cultures were then incubated for 6h (to allow time for OT-1 cells to become activated and produce IFNγ), fixed, and stained for pSTAT1 (Fig 3B). We observed spherically symmetric, spatially-confined niches of pSTAT1 surrounding OT-1 T cells (Fig 3C-D, S3B). Untreated cultures demonstrated no pSTAT1 staining, while cultures treated with a saturating dose of IFNγ demonstrated widespread, uniform STAT1 activation (Fig S3A).

**Figure 3:**
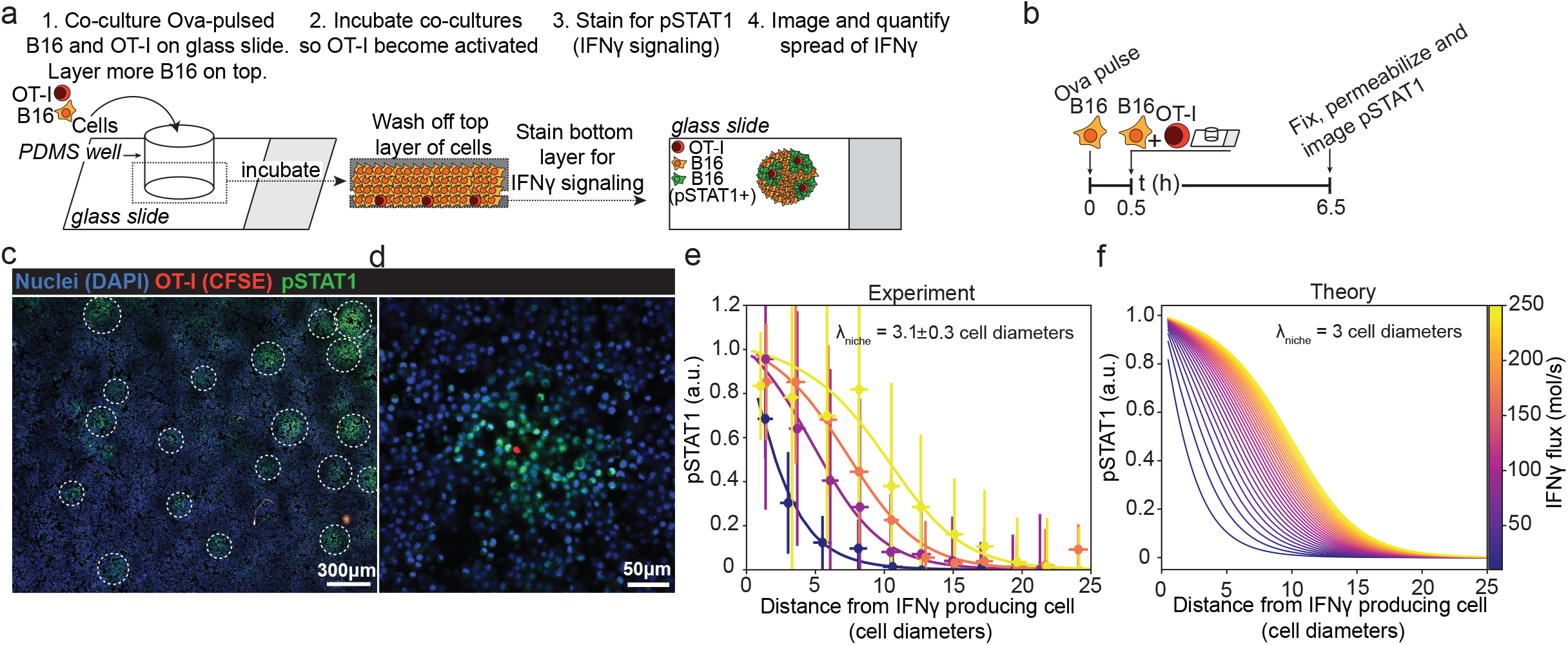
Imaging reveals niches of IFNγ signaling surrounding activated T cells, and enables direct measurement of the signaling length scale. **(a)** Schematic demonstrating the principle of the PlaneView technique that enables dense, 3-D cell culture and convenient 2-D imaging. **(b)** Experimental diagram of B16-OT-1 T cell PlaneView co-culture experiment. **(c)** Zoomed out view of clusters of pSTAT1 staining in B16 cells after co-culture with OT-1 T cells. **(d)** Close-up image of a cluster of pSTAT1 staining in B16 cells surrounding one OT-1 T cell. **(e)** Example quantifications in the decay in pSTAT1 staining surrounding OT-1 T cells. Data is shown as points with error bars and solid lines depict fits from our theoretical diffusion-consumption equation. **(f)** Theory depicting expected pSTAT1 decay profiles given a defined length scale of 3 cell diameters and varying the IFNγ flux (*a*.*k*.*a*. IFNγ production rate by T cells).

We used a theoretical framework based on Diffusion-Consumption dynamics to directly quantify the length scale (λ_*niche*_) of cytokine spread from these imaging experiments ^31,49^. In our framework, cytokines originate from a source, diffuse freely between cells, eventually bind to a receptor, and are endocytosed. The amount of consumed cytokine linearly depends on the density of consuming cells. Consuming cells that are closer to the production source have a higher probability of capturing and depleting cytokines than cells further away, which creates a gradient in the concentration profile of the cytokine. The mean time (*τ* _*diffusion*_) that IFNγ diffuses before being captured is inversely proportional to the density of IFNγ consuming cells (*n* _*consumers*_) multiplied by its kinetic rate of consumption (*k*) . A molecule of IFNγ would therefore travel an average distance of 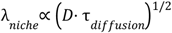, where *D* is the molecular diffusion coefficient. The signaling length scale is then expected to be 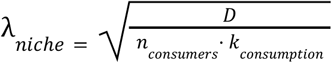.

The full profile of pSTAT1 depends on the signaling length scale λ _*niche*_, which is the rate of decay of cytokine concentrations in the sub-saturating regime, and the rate of cytokine production by the producer which scales the extent of the saturating regime (Fig 3F, S3D-F). We expect λ _*niche*_ to be global across the entire sample while the production rate, and therefore length of the saturation regime, varies between different producers. We globally fit the observed pSTAT1 profiles to the theoretically expected expression ^31^ with excellent agreement (Fig 3E, S3C). We obtained a direct measurement for IFNγ’s characteristic length scale: 3.1 ± 0.3 cell diameters, or approximately 30μm. As predicted by theoretical considerations, the observed curves are fit with a global length scale, while differences in production rates between producers generate variability in the saturating regime of pSTAT1 profiles (represented by different colors in Fig 3E-F, S3D).

These data demonstrate how a diffusion-consumption mechanism spatially confines IFNγ spread around cytokine producers, leading to compact niches of high cytokine concentration. We therefore expect the density of cytokine producers to be important for determining variability in IFNγ signaling and gene expression within a tumor.

### Simulation and theory show a transition from isolated niches to overlapping fields with increasing cytokine producer cell density

To determine how varying the density of IFNγ-producing cells affects the distribution of single-cell responses throughout the tumor, we turned to simulation and theory. For simulations, cells were assumed to lie on a 3-D cubic lattice, and a varying fraction of randomly selected cells were assigned to be cytokine producers. Individual producers generated a spherically symmetric cytokine niche around themselves, corresponding to a constant IFNγ flux of 50 mol/s per T cell, and a λ_*niche*_ = 30μm, as measured in Fig 3E and S3C (Fig 4A). Starting with a very small fraction of cytokine producers, we observe isolated niches of cytokine response (Fig 4A). As the density of producers increases, these niches begin to overlap, creating a more homogenous population of responding cells.

**Figure 4:**
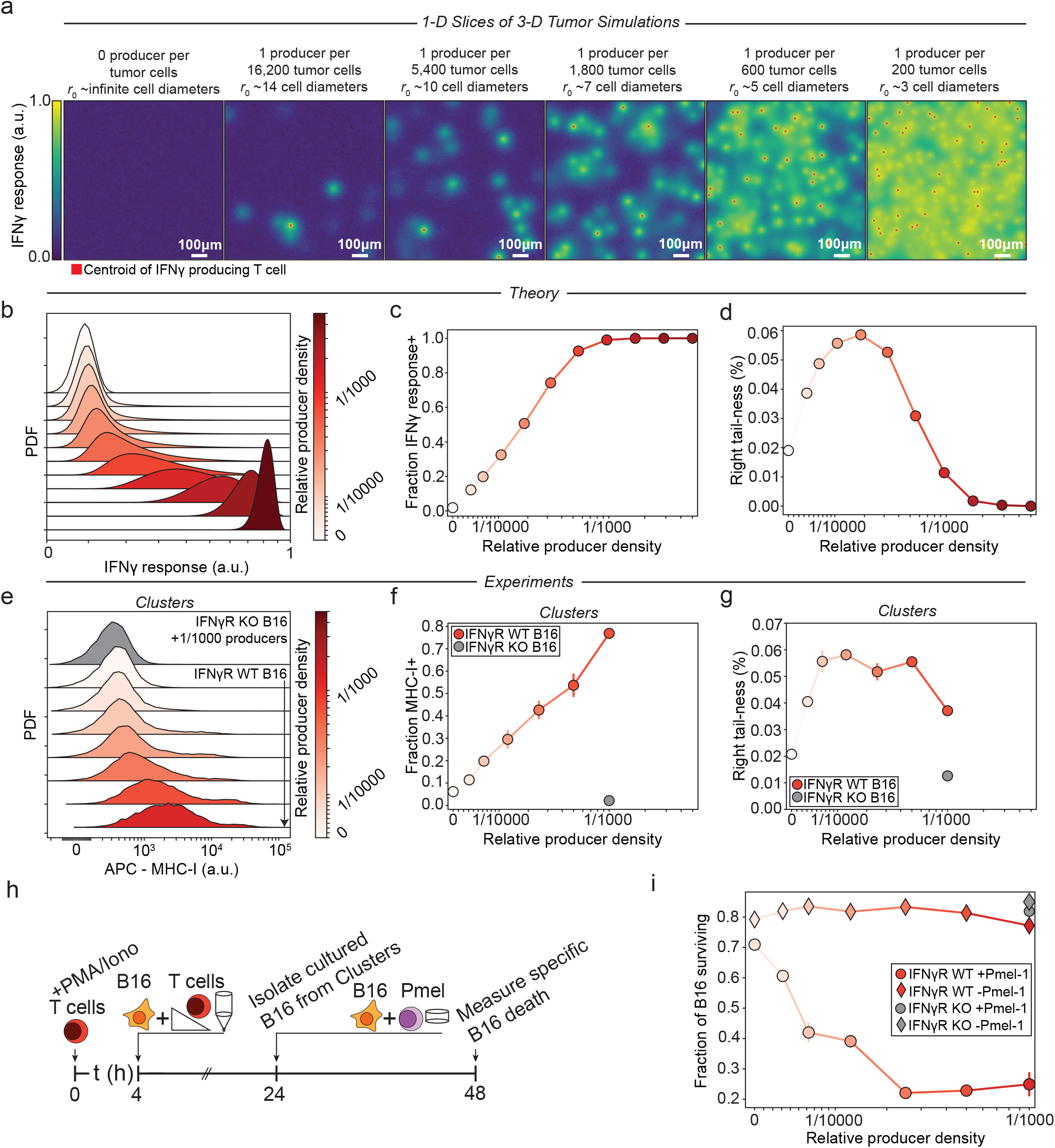
Gradients of IFNγ translate into variability in antigen presentation and susceptibility to T cell mediated killing. **(a)** Representations of 1-D slices of 3-D simulations. For simulations, cells were embedded in a 3-D cubic lattice, and a varying fraction of randomly-selected cells were assigned as cytokine producers. Individual producers generated a spherically symmetric cytokine niche around themselves, corresponding to a constant IFNγ flux of 50 mol/s per T cell, and a λ_*niche*_ = 30μm, as measured in our experiments. Red squares represent the centroid of a cytokine producer. Squares are not always present in the center of a cytokine response niche because, in many cases, the producer centroid lies outside of the 1-D slice. **(b)** Histograms of simulated IFNγ response of cells interspersed randomly with varied densities of IFNγ-producer cells. **(c)** Simulation results for the fraction of cells responding to IFNγ as a function of varied IFNγ producer density. **(d)** Simulation results for the right-tailness (>2 standard deviations above the population mean) of cellular response to IFNγ as a function of varied IFNγ producer density. **(e)** Histograms of MHC-I levels on B16 cells co-cultured with activated T cells. T cells were stimulated with PMA and Ionomycin for 4h before being co-cultured at varying densities with B16 cells overnight in clusterwells. Cells were then stained for MHC-I (H2-K^b^) and measured by flow cytometry. **(f)** Quantification of the fraction of MHC-I+ cells from (e). **(g)** Quantification of the right-tailness (>2 standard deviations above the population mean) of MHC-I levels from (e). **(h)** Experiment diagram for results shown in (i). T cells were stimulated with PMA and Ionomycin for 4h before being co-cultured at varying densities with B16 cells in clusterwells overnight. Cells were then collected from clusterwells and co-cultured with Pmel-1 CD8+ T cells. Pmel-1 mediated killing was quantified by flow cytometry analysis of DAPI incorporation by B16s. **(i)** Quantification of the fraction of surviving B16s as a function of the density of co-cultured IFNγ-producing T cells.

We next aimed to theoretically predict the distribution of cytokine responses in the population. We assume that (at least locally) cytokine producers are randomly distributed. We then calculate the concentration of cytokine at each position in space by summing over the cytokine concentrations generated by all the producers. The calculation is similar in principle to that employed for fluorescence intensity-distribution analysis ^50^, and results in a distribution of cytokine exposure of the tumor cells. A full description of the approach is provided in the supplementary materials.

At the population level, heterogeneity in IFNγ exposure translates into broad, long-tailed distributions of IFNγ response (Fig 4B). At high densities of cytokine producers, the system enters into a saturation regime where most cells are exposed to IFNγ, ultimately leading to a population-wide shift in IFNγ response (Fig 4B-C). To quantify the cells exposed to high concentrations of IFNγ, we computed the proportion of cells with an IFNγ response greater than two standard deviations above the mean, a metric we term “right-tailness”. A normal distribution has a right-tailness of ∼0.02, and a value significantly greater than that indicates a distribution with a heavy right-tail. Right-tailness initially increases along with the density of cytokine producers, then peaks and decreases at higher producer density (Fig 4D). The decrease at high cytokine producer densities reflects the shift into saturation as the entire population responds to IFNγ.

The distributions computed from our theoretical framework are strongly concordant with those generated from our simulations (Fig 4B, S4A-C). Taken together, our simulations and theory argue that at low to medium densities of IFNγ-producing cells, IFNγ is only present in compact domains or niches, generating cellular variability in IFNγ response. Only in situations where the density of IFNγ-producing cells is high do we expect a global response to IFNγ due to overlap of niches.

### Steep gradients of IFNγ translate into variability in antigen presentation and susceptibility to T cell mediated killing

To test the validity of our theoretical results, we set out to measure gene expression and T cell killing in an *in vitro* tumor model with precisely controlled ratios of TILs to tumor cells. We selected the antigen presentation pathway as a measure of IFNγ-responsive gene expression ^17^. To measure tumor expression of MHC Class I (MHC-I), we co-cultured activated, IFNγ-producing T cells at varying densities with B16 cells in clusterwells. A uniform population of IFNγ-producing T cells was generated as described above by activating the cells for several hours with PMA and Ionomycin (Fig S2E-F).

After overnight co-culture, we observed broad, long-tailed distributions of MHC-I expression on B16 cells that depended on the density of IFNγ producing cells (Fig 4E-F). These broad, long-tailed distributions are in agreement with the results of our simulations and theory (Fig 4B-C, S4A). In contrast, B16 cells exposed to a dose titration of IFNγ in well-mixed conditions exhibit a narrow, uniform upregulation of MHC-I similar to pSTAT1 (Fig S4D-E, S2A). These data indicate that, in the clusterwells, B16 cells are exposed to varying concentrations of IFNγ and therefore show cell-to-cell variability in MHC-I expression.

Consistent with our simulations, we observe that right-tailness increases dose-dependently with the density of T cells, peaks, and then decreases at T cell densities greater than about 1 in 8000 cells (Fig 4D and G). We also calculated skewness, a measure of the asymmetry of a probability distribution, and observe a characteristic heavy right tail (positive skewness) at low producer densities, followed by a left tail (negative skewness) near saturation (Fig S4G-I). These measures contrast starkly with those obtained from well-mixed cultures, where right-tailness decreases consistently with increasing IFNγ concentration (Fig S4F) and never exceeds the levels expected for a normal distribution.

Finally, we evaluated the functional effect of MHC-I upregulation on T cell mediated killing. We co-cultured activated, IFNγ-producing T cells at varying densities with B16 cells in the clusterwells (Fig 4H). After overnight co-culture, the cells were removed from clusterwells and co-cultured with CD8+ cytotoxic Pmel-1 T cells, which bear a B16 tumor antigen (gp100)-specific transgenic TCR ^51^. An excess of IFNγ-neutralizing antibodies was added to eliminate any effect of IFNγ-production arising from Pmel-1 activation. We measured Pmel-mediated killing by quantifying DAPI incorporation by B16 cells, and observed a sharp drop-off in the proportion of surviving B16s according to the initial number of co-cultured IFNγ-producing T cells (Fig 4I). This killing was Pmel-1-specific and required IFNγ sensing by B16 cells as IFNγR KO B16s were not killed in any context. We previously showed that blocking MHC-TCR interaction during IFNγ stimulation abrogates Pmel-mediated killing, demonstrating that the pro-death effect of IFNγ depends on MHC-I upregulation ^17^.

These results demonstrate that the variable upregulation of MHC-I on B16 cells achieved in dense co-cultures translates into a functional difference in susceptibility to T cell killing. Notably, Pmel-1 killing saturates only after the entire population of B16s begins to up-regulate MHC-I, indicating that B16 cells that are outside the niches where IFNγ concentrations are high can escape T cell-mediated killing. These results argue that widespread IFNγ signaling throughout the entire tumor is critical for immune regression.

### Human melanomas exhibit niches of IFNγ signaling that surround TILs

To examine the spatial pattern of IFNγ signaling in human melanoma, we analyzed a patient-derived FFPE primary cutaneous melanoma lesion that was multiplex antibody stained for a panel of tumor, signaling, and immune markers using CyCIF ^52^. We first took an unbiased look at the spatial relationship between each stained marker in the tumor by computing Moran’s I, a global measure of spatial correlation between pairs of markers, then performing hierarchical clustering to identify groups of correlated markers (Fig 5A) ^53,54^.

**Figure 5:**
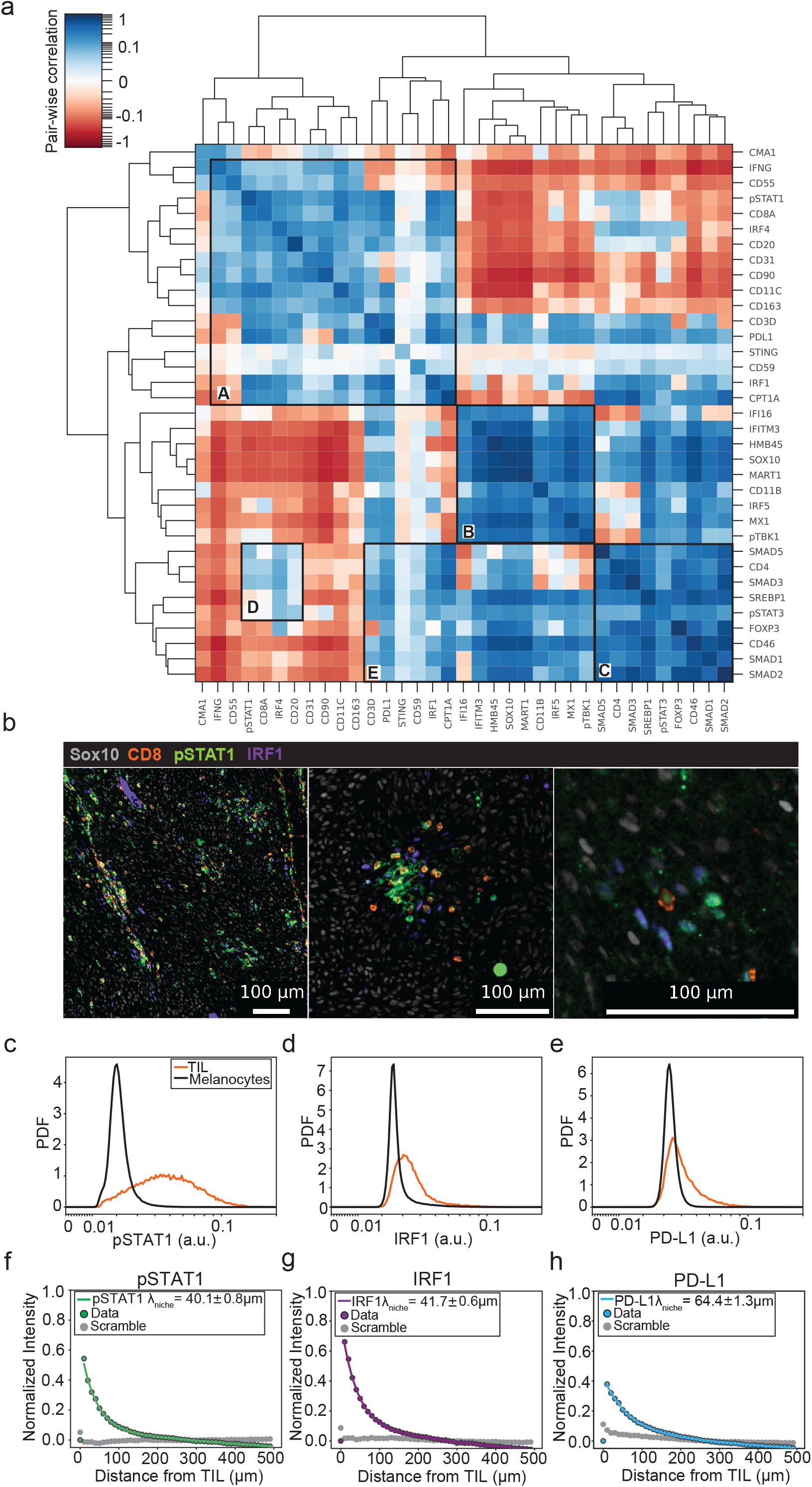
Human melanomas exhibit spatially confined niches of IFNγ signaling that surround TILs. Clustergram of Moran’s I values computed from human primary cutaneous melanoma. Moran’s I is a global measure of spatial correlation between pairs of stained markers in adjacent cells. Computed values were then clustered hierarchically, and groups of correlated markers were annotated manually. Images of regions of human tumor depicting the spatial pattern of IFNγ signaling through pSTAT1 and IRF1 staining, surrounding CD8+ T cells. **(c-e)** Probability distribution functions for pSTAT1 **(c)**, IRF1 **(d)**, and PD-L1 **(e)** in TILs and Sox10+ melanocytes. **(f-h)** Decay in the mean normalized intensities of pSTAT1 **(f)**, IRF1 **(g)**, and PD-L1 **(h)** as a function of distance from TILs. Data are shown as blue and orange points with error bars, and the solid green line depicts the fit from our theoretical diffusion-consumption equation.

This analysis revealed several groups of spatially-correlated and anti-correlated markers, which we annotated as clusters A-E. We focus here on groups of correlated markers, although the anti-correlated markers merit further investigation. We refer to cluster A, which is enriched with markers for activated CD8 T cells, B cells, antigen presenting cells, leukocyte adhesion and migration markers, and IFNγ signaling, as “Mature Adaptive Immunity”. Cluster B, “Innate Immunity”, is enriched with markers of type I IFN signaling, antiviral signaling, tumor antigens, and macrophages. Cluster C, “Immunosuppressive”, contains primarily markers of TGFβ signaling and regulatory T cells (Tregs). Cluster D, “Nascent Adaptive Immunity”, contains evidence of active CD8 T cells, but not other immune cell subsets. Finally, Cluster E, “Myeloid Suppressive Niche”, is enriched with tumor antigens, IFNγ signaling, macrophages, innate immune signaling, and Tregs. We were particularly interested in clusters A and E, because they demonstrate a strong, positive spatial relationship between activated T cells, IFNγ, and markers of IFNγ signaling and downstream gene expression, such as pSTAT1, IRF1, and PD-L1. To investigate this relationship further, we visually examined tumor images.

Images demonstrate niches of IFNγ signaling, as evidenced by pSTAT1 and IRF1 staining, that surround individual TILs or groups of TILs (Fig 5B). Often, regions of the tumor that are only a few cell diameters away from TILs exhibit no IFNγ signaling. This indicates that there are steep gradients of IFNγ present in the tumor: in close proximity to activated TILs, IFNγ is present at high concentrations, whereas it is essentially non-existent further away.

To quantify the prevalence of IFNγ signaling throughout the lesion, we calculated distributions of IFNγ-responsive signaling (pSTAT1) and gene expression (IRF1, PD-L1) for different cell subsets within the tumor. Melanocytes (tumor, atypical, and typical) were defined by expression of Sox10 ^52^. TILs were defined by their expression of CD3, CD4 or CD8, CD45RO, absence of FoxP3, and spatial localization of less than 50μm from at least 8 tumor cells (see Fig 1). Only a very small proportion of total tumor cells exhibit staining for IFNγ signaling or gene expression (Fig 5C-E). By contrast, a high proportion of TILs show a response to IFNγ, unsurprisingly since TILs are the main source of IFNγ ^27^ and are exposed to it via autocrine signaling.

We next computed the spatial cross-correlation of TILs with the mean normalized intensities of pSTAT1, IRF1, and PD-L1 (Fig 5F-H). As a control, we used values of pSTAT1, IRF1, or PD-L1 intensities from sites sampled randomly throughout the tumor (“scrambled”). These data demonstrated a steep drop off in IFNγ signaling and gene expression as a function of distance from TILs. We fit these cross-correlation profiles to the theoretically expected expression and measured λ _*niche*_ = ∼40μm for both pSTAT1 and IRF1, the immediate signaling responses to IFNγ, and ∼65μm for PD-L1 which is further downstream. We hypothesize that the longer length scale measured for PD-L1 is due to cellular migration, which happens over slower timescales than signaling. These values are comparable to those measured directly in mouse B16 cells (∼30μm) (Fig 3E), and indicate that IFNγ spread is highly spatially confined in human tumors as well as mouse melanoma cells.

Recently, another group analyzed single cell RNA-sequencing data and drew an opposite conclusion to ours: that IFNγ spreads widely throughout human melanomas ^27^. However, due to technical challenges, this work largely analyzed the IFNγ gene expression response signature in monocytes, macrophages, and neutrophils. The interpretation of wide IFNγ spread relies on the assumption that such cells are evenly spaced throughout the tumor. We find that this is not the case - monocytes, macrophages, and neutrophils are highly spatially correlated with TIL-rich regions of the tumor (Fig S5A-C). Accordingly, tumor-infiltrated macrophages exhibit much higher pSTAT1, IRF1, and PD-L1 staining than tumor cells (Fig 5SD-F). We therefore conclude that IFNγ is highly spatially confined to regions of local T cell activation and anti-tumor activity.

## Discussion

IFNγ is a critical anti-tumor cytokine that is required for checkpoint blockade therapies to work. It is therefore important to quantify the extent to which IFNγ spreads in the tumor microenvironment, as well as the factors that govern that spread. Past work has yielded contradictory results that span a wide range: from essentially nearest neighbor signaling, to perfusion of the cytokine over several hundred microns. We applied a quantitative framework based on diffusion and consumption dynamics to study the spread of IFNγ *in silico*, in dense 3-D *in vitro* conditions, and in human tumors. We observed that, at least in melanomas, IFNγ signaling is highly spatially confined to regions in the immediate vicinity of activated IFNγ-producing T cells, and decays with a characteristic length scale of 30-40μm or ∼3-4 cell diameters. Similar results were observed in dense, 3D, *in vitro* settings and in clinical samples. Spatial constriction of IFNγ spread generated significant cell-to-cell variability in STAT1 signaling, antigen presentation, and susceptibility to T cell-mediated killing. Further, our data and simulations suggest that widespread IFNγ signaling is only possible when the density of IFNγ-producing TILs is high enough for localized cytokine fields to overlap.

An important aspect of our work is its generalizability and ability to account for different measurements of IFNγ spread. We measured a spreading length scale of ∼30-40μm for IFNγ in mouse B16 cells and human melanomas. Because the length scale is determined by diffusion and consumption, it can be tuned by changes in the parameters that determine consumption, such as receptor expression or consumer cell density. For example, we previously demonstrated variation of ∼50% (from 80-120μm) in the length scale of Interleukin-2 spread in the lymph nodes, depending on the density and receptor-expression of IL2-consuming regulatory T cells. We argue that the same is true for IFNγ - highly confined spread and slightly longer-distance spread are both plausible depending on IFNγ consumption in the tumor.

It is important to consider the potential limitations of comparing data from human, mouse, and *in vitro* studies. *In vitro* and mouse studies often use TCR-transgenic systems or drug/agonist-antibody-based T cell activation, which results in a high level of T cell activation and an artificially high number of tumor-reactive T cells. This leads to a synchronized wave of T cells producing IFNγ in the tumor, which may not accurately reflect the proportion of T cells producing IFNγ in human tumors (estimated at around 10% of total T cells ^27^). As a result, the spread of IFNγ in natural settings may be overestimated or misinterpreted.

One of the most commonly-accepted positive prognostic factors for tumor regression is the number of TILs ^55^. Put simply: more is better. We argue that in addition to the number of TILs, the spatial distribution of these cells within the tumor is also important. A high density of TILs concentrated in one region of the tumor may not be as effective as a lower density of TILs that are evenly distributed throughout the tumor. Indeed, newer prognostic factors incorporate more refined and complex tumor-immune measures, such as average distance between tumor cells and TILs ^56^ and localization of specific TIL subsets in key regions of the tumor ^57^. Finally, solid tumors have been analogized as collections of cellular “neighborhoods” ^52,58–60^. One neighborhood may exhibit inflammatory regression, while only a few cell diameters away, another neighborhood is immunosuppressive. This work supports these ideas by providing a mechanism that can spatially confine immune molecules to small regions within a tumor, thereby generating non-genetic cell-to-cell variability.

## Methods

### Mice

C57Bl/6J and Pmel-1 CD8+ TCR Transgenic mice were purchased from The Jackson Laboratory, and OT-1 mice were provided by Dr. Arlene Sharpe (Harvard Medical School). All mice were maintained in SPF conditions at an Association for Assessment and Accreditation of Laboratory Animal Care-accredited animal facilities (Harvard Medical School).

### Human tumor specimens and CyCIF

Human tumor images used in this paper were originally selected, processed, cataloged and described by D. Liu and J-R. Lin ^44^ and A. Nirmal and Z. Maliga ^52^. In each case, samples were collected under Institutional Review Board (IRB) approval by either the Brigham and Women’s Hospital IRB (Protocol #18-1363) ^52^ or the Dana-Farber/Harvard Cancer Center IRB (Protocol #11-181) ^44^. CyCIF was performed as described in ^44–46,52^ and at protocols.io (https://dx.doi.org/10.17504/protocols.io.bjiukkew). Access to raw imaging data was provided by the Laboratory of Systems Pharmacology at Harvard Medical School.

**Table 1.**
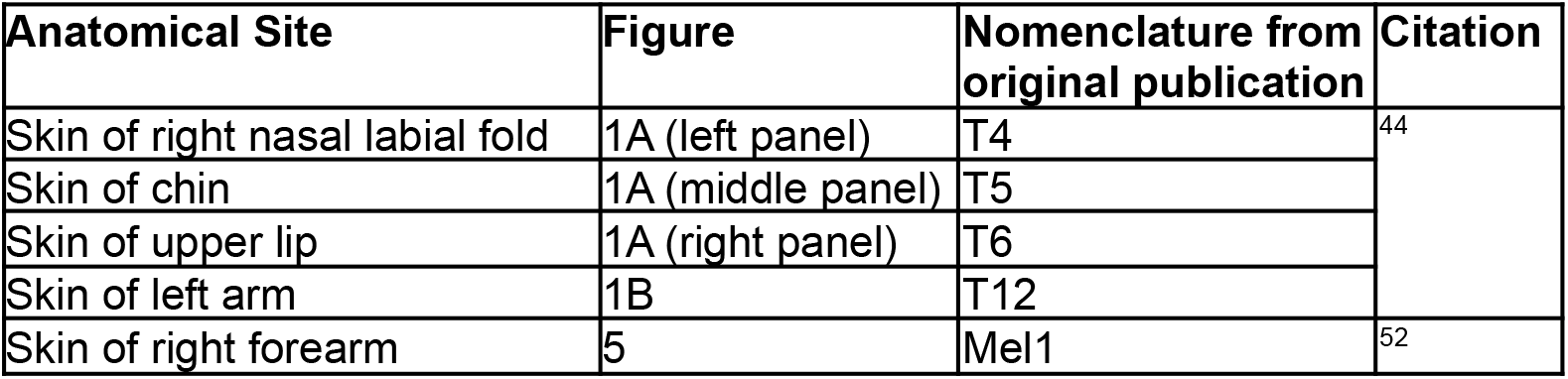
Human tumor specimens shown in Figs. 1 and 5.

| Anatomical Site | Figure | Nomenclature from original publication | Citation |
| --- | --- | --- | --- |
| Skin of right nasal labial fold | 1A (left panel) | T4 | <sup>44</sup> |
| Skin of chin | 1A (middle panel) | T5 |  |
| Skin of upper lip | 1A (right panel) | T6 |  |
| Skin of left arm | 1B | T12 |  |
| Skin of right forearm | 5 | Mel1 | <sup>52</sup> |

### Cells and Culture Conditions

B16-F10 cells were purchased from ATCC (#CRL-6475) or were provided by Dr. Grégoire Altan-Bonnet (National Institutes of Health) and maintained in complete culture media: RPMI supplemented with 10% heat inactivated fetal calf serum (Sigma, US Origin), 2mM L-glutamine, 10mM HEPES, 0.1mM non-essential amino acids, 1mM sodium pyruvate, 100μg/ml of penicillin, 100μg/ml of streptomycin, and 50μM β-mercaptoethanol. Primary T cells were harvested from the lymph nodes and spleen, activated using 10ng/ml PMA and 500ng/ml Ionomycin, and cultured for 3 days. Dead cells were removed using Ficoll-paque plus (GE Healthcare) and subsequently cultured in complete culture media supplemented with 5 nM human IL-2 (Peprotech) for up to 10 days. Where indicated, IFNγ was induced by restimulating T cells with 10ng/ml PMA and 500ng/ml Ionomycin for 4–6 hours. To differentiate OT-1 or Pmel-1 CD8+ T cells before co-culturing, splenocytes were activated with 10ng/ml PMA and 500ng/ml Ionomycin for 2 days, then ficolled and cultured for an additional 3-6 days in 2nM IL2.

Where indicated, between 0.5-2×10^5^ B16 cells were seeded into either 96 well plates or sterilized Clusterwell plates in 200μl of total media, and centrifuged at 400 rcf for 1m to sediment cells. Pelleted cells were then stimulated with the indicated concentration of IFNγ pipetted gently into the media on top, taking care to avoid disturbing the cell pellet.

For B16-IFNγ-producing T cell co-cultures, we used OT-1 CD8+ T cells, which are expected to have no reactivity towards B16 cells in the absence of OVA. OT-1 cells were induced to produce IFNγ using PMA and Ionomycin for several hours, then co-cultured with B16 cells in either 96 well plates or sterilized Clusterwell plates at varied T cell density, keeping the total cell number at 2.5×10^5^ cells per well.

For Pmel-1 killing assays, IFNγ-producing OT-1 and B16 cells were co-cultured at the indicated densities overnight in Clusterwells. The next day, the cluster of cells was dispersed and re-cultured in 96 well plates at a 1:1 ratio with differentiated Pmel-1 CD8+ T cells for 8 hours, keeping the total number of cells at 2×10^5^ cells per well. To neutralize effects of any residual IFNγ produced by activated Pmel-1 cells, anti-IFNγ (XMG1.2: Tonbo) was supplemented at 2μg/ml. B16 death was quantified by DAPI incorporation.

### PlaneView Imaging Experiments

Experiments were performed as previously described, with some small modifications. First, glass slides were coated with poly-l-lysine for 40m at 37°C. Next, a hollow cylinder of PDMS (with 6mm diameter) was placed on the slide. PDMS rapidly attaches to the slide, creating a small well. Prior to loading into wells, B16 cells were pulsed for 0.5h with 5μg/ml OVA_257-264_ (Genscript), and then washed twice in RPMI. Differentiated OT-1 T cells were then labeled with 1μM carboxyfluorescein succinimidyl ester CFSE (Molecular Probes) for subsequent identification. To create a densely-packed 3D cell patch, cell mixtures are added to the PDMS wells and the slide is then centrifuged at 800 rcf for 1m. This step is undertaken in a two step manner. First, OVA-pulsed B16 cells are mixed with 0.05% OT-1 T cells, deposited in the PDMS well, and centrifuged to form a monolayer. Next, approximately 10 layers of OVA-pulsed B16 cells alone (*i*.*e*. no OT-1 cells) were deposited on top. This creates a dense, 3D system where IFNγ production only originates from T cells located in the bottom-most layer, thereby simplifying our interpretation of images and quantification of IFNγ spread. As negative and positive controls, we prepared patches of B16 cells alone or B16 cells supplemented with 10nM of IFNγ. Cultures are incubated for 1h at 37°C, allowing them to reach steady state, and then fixed with warm 4% PFA for 15m and permeabilized with ice cold 90% methanol for at least 30m at -20°C.

At this point, the patch of cells can be analyzed using conventional immunofluorescence. After permeabilization, the top layers of B16 cells wash off, and we are left with a monolayer of OT-I T cells mixed with B16 cells. Non-specific antibody binding is prevented by first blocking for 1h in 5% FCS and 0.3% Triton-X-100. Primary antibodies (anti-pSTAT1, clone 58D6, Cell Signaling Technologies) are applied at a dilution of 1:200 in blocking buffer in moist chambers for 1h at room temperature. After washing, secondary antibodies (anti-Rabbit Alexa 647, #711-606-152, Jackson ImmunoResearch) were then applied at a dilution of 1:300 as already described. Patches were then briefly stained with DAPI and coverslipped using Fluoromount Aqueous Mounting Media.

### Flow Cytometry

T cells were fixed for 10 min in 1.6% PFA on ice, and subsequently permeabilized in 90% methanol for at least 30 min at −20°C. Cells were stained with anti-CD4 (clone RM4–5, Tonbo), anti-CD8α (clone 53-6.7, Tonbo). Where indicated, T cell IFNγ production was quantified using the mouse IFNγ secretion kit (Miltenyi Biotech). B16-F10 cells were fixed as above and stained with anti-pSTAT1 (58D6: Cell Signaling Technologies). Live B16-F10 cells were stained with anti-MHC-I H2-K^b^ (AF6-88.5: Biolegend).

Flow cytometric data were collected on an LSR II (BD Biosciences). Analysis was done with FlowJo software (TreeStar), and with custom code written in python and available on GitHub (https://github.com/oylab/oyFlow).

### Confocal Microscopy

Confocal microscopy of PlaneView specimens was performed on a Zeiss LSM510META laser scanning confocal microscope equipped with lasers emitting 458, 488, 514, 565 and 633nm.

### Image Analysis

Automated image processing and analysis was performed using custom code written in Python. Software is available from the GitHub repository: https://github.com/oylab/oyLabImaging Cell nuclei were segmented using cellpose nuclei model ^61^ based on the nuclear dye DAPI or Hoechst 33258 with diameter 5 for PlaneView images (Fig 3) and 20 for CyCIF images (Figs 1 and 5). The single-cell position and multi-channel intensities were then extracted and stored.

## Statistical Analysis and Data Presentation

All relevant data are shown as mean ± standard error of the mean (s.e.m.) or standard deviation (s.d.). Statistical tests were selected based on appropriate assumptions with respect to data distributions and variance characteristics.

## Author Contributions

Conceptualization, J.O.-Y. and A.O.-Y. Investigation, J.O.-Y., A.O.-Y., E.C., C.W., S.I., and O.K. Software, A.O.-Y. Writing, J.O.-Y. and A.O.-Y. Resources, J.O.-Y.

## Acknowledgements

We thank Drs. Peter K. Sorger, Ajit Nirmal, and Jia-Ren Lin for providing and facilitating access to human tumor microscopy data; Arlene Sharpe and Grégoire Altan-Bonnet for sharing mice and cell lines, Kathy Buhl and Samantha Jalbert for laboratory support, and Dr. Rebecca Ward for critically reading the manuscript and providing feedback.

## Declaration of Interests

The authors declare no competing interests.

## Notes

### Competing Interest Statement

The authors have declared no competing interest.

