## Supplemental Materials for "The Spread of Interferon-γ in Melanomas is Highly Spatially Confined, Driving Non-Genetic Variability in Tumor Cells"

Contents:

1. Supplemental figures
2. Mathematical framework
    a. Cytokine concentration and response around single producer
    b. Response distribution in dense tissues

1. Supplementary figures

*Supplement to Figure 1: Demonstration of relevant single CyCIF image channels and exclusion of non-infiltrated T cells.*

**(a)** Region of high density of TILs from tumor site 2. Individual channels for relevant markers are shown to demonstrate staining quality. **(b)** Image reconstructions depicting TILs and Non-TILs from tumor site 2. Non-TILs express the relevant markers to be assigned as T cells, but are not located within at least 25µm of at least 10 S100A+ tumor cells.

### Supplement to figure 1

a

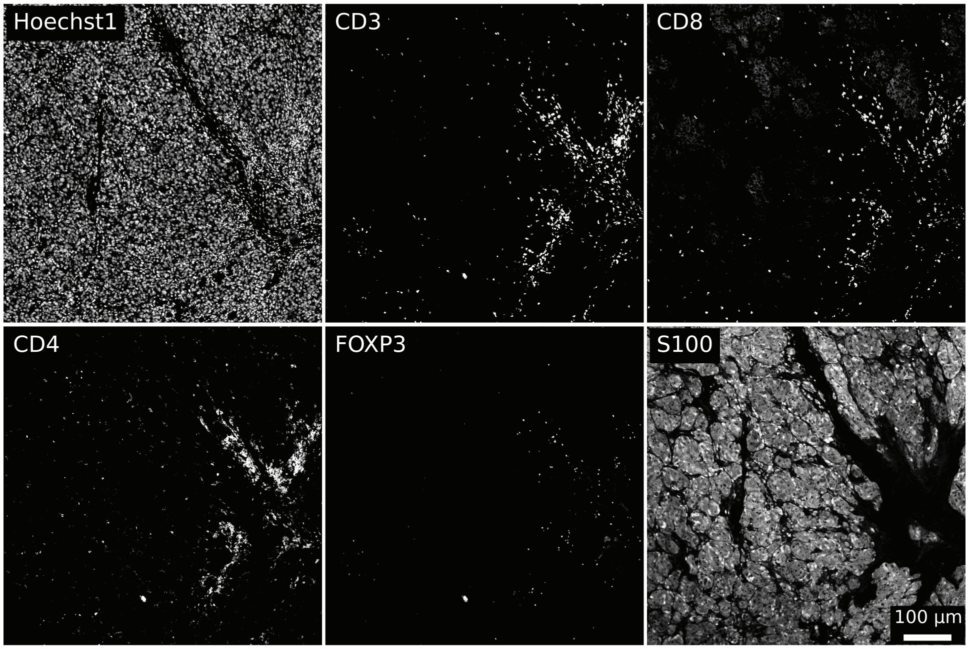


b

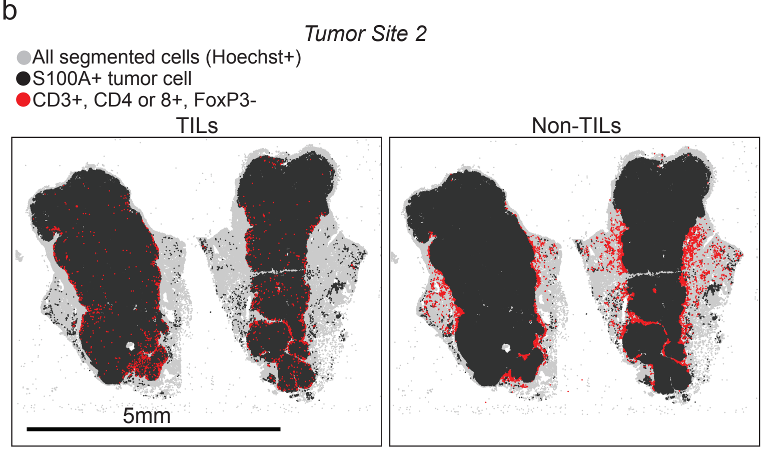

*Supplement to Figure 2: Validation of B16 signaling responses to IFNγ and quantification of IFNγ production in PMA and Ionomycin activated T cells.*

**(a)** Histograms of pSTAT1 levels in IFNγ-stimulated B16 cells cultured in conventional 96 well plates. **(b)** Normalized geometric mean fluorescence intensities of pSTAT1 levels from (a). **(c)** Histograms of pSTAT1 levels in IFNγ-stimulated B16s cultured in either 96 well plates or clusterwells for 90m (as opposed to 60m in Figure 2). **(d)** Percentages of pSTAT1+ cells from (c). **(e)** Histogram of IFNγ intracellular staining in PMA and Ionomycin activated T cells. Brefeldin A was added during the last 2 hours of the total 4h PMA and Ionomycin stimulation, and IFNγ production was quantified by flow cytometry. **(f)** Quantification of the percentage of IFNγ+ cells from (e).

### Supplement to figure 2

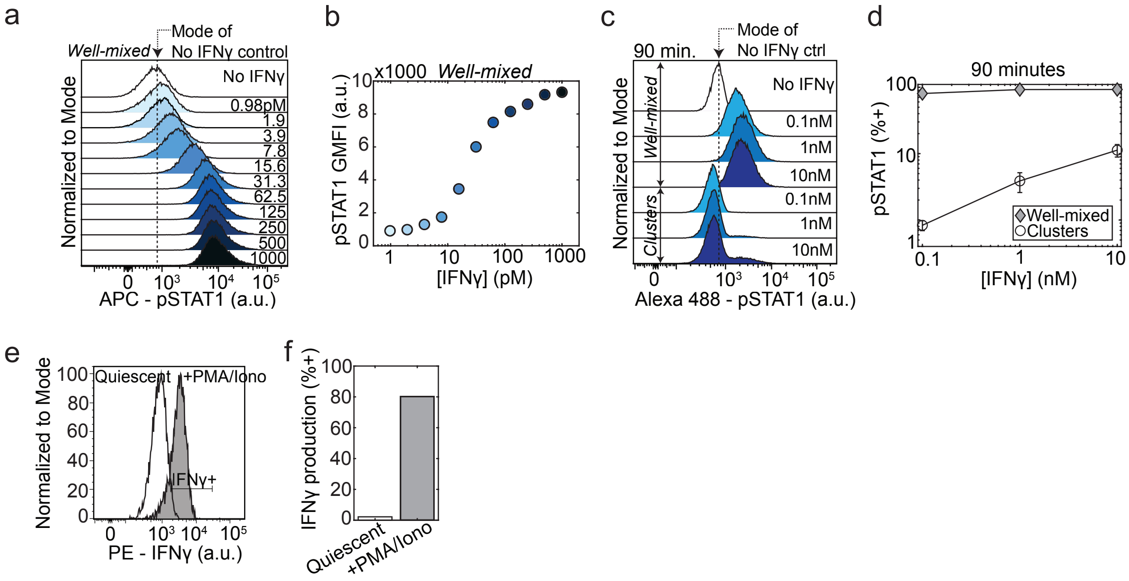

*Supplement to Figure 3: Validation of PlaneView imaging and visualization, demonstration of fits from additional pSTAT1 signaling zones.*

**(a)** Zoomed out images showing the background level of pSTAT1 staining in B16 cells in the absence of T cells or after culture with a saturating dose of IFNγ. **(b)** Examples of additional close-up images of pSTAT1 staining in B16 cells surrounding one OT-1 T cell. **(c)** Examples of many additional pSTAT1 clusters globally fit to our theoretical diffusion-consumption equation. **(d)** Theory depicting expected IFNγ decay profiles given a defined length scale of 3 cell diameters and varying the IFNγ flux (*a.k.a.* IFNγ production rate by T cells). **(e)** Theory depicting expected pSTAT1 signaling $EC_{50}$ values given a defined value for IFNγ flux and varying the length scale of spread. **(f)** Theory depicting expected pSTAT1 levels given a defined value for IFNγ flux and varying the length scale of spread.

### Supplement to figure 3

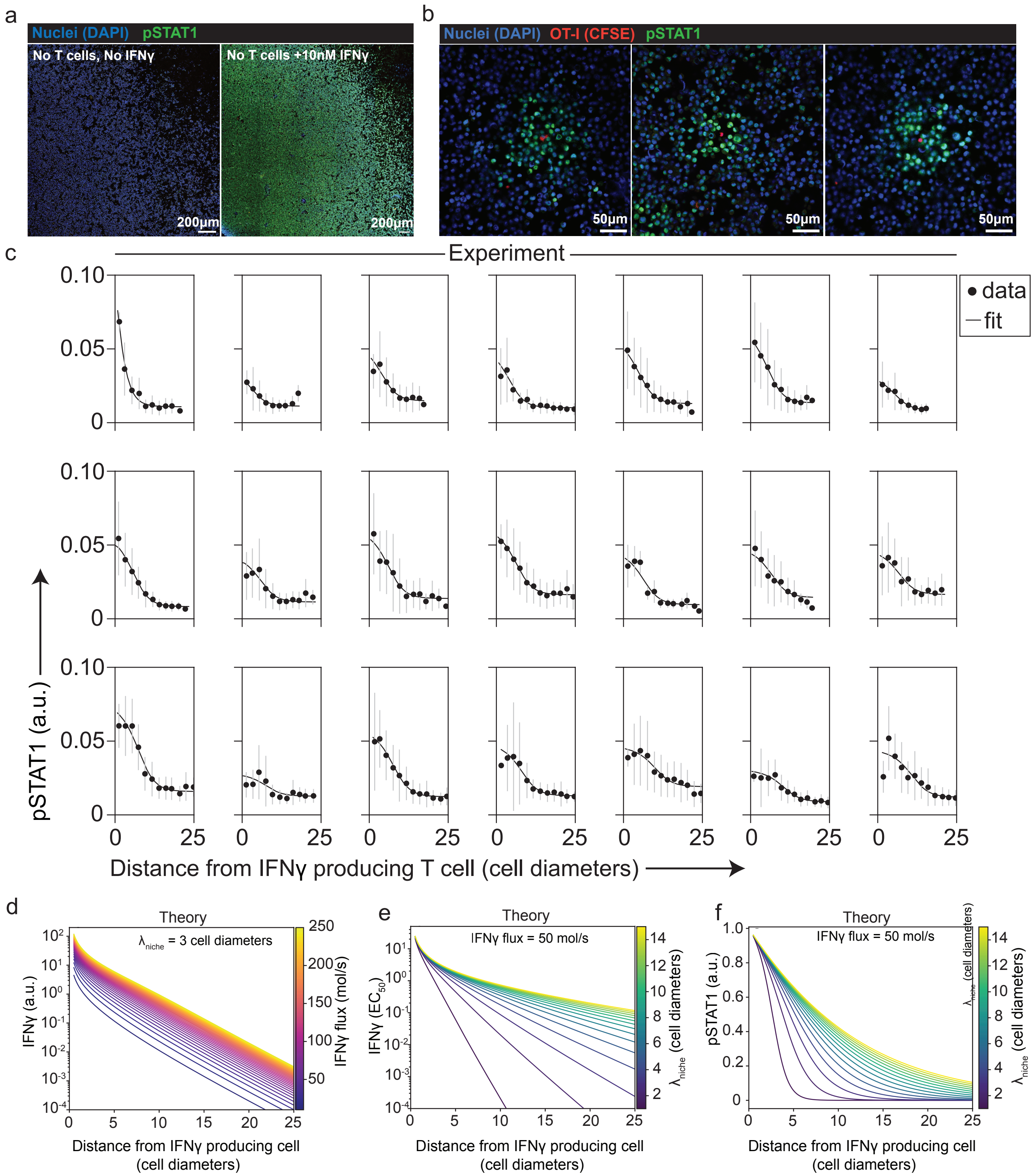

*Supplement to Figure 4: Comparison of theory and simulations, and validation of B16 antigen presentation response to IFNγ.*

**(a)** Theoretical distributions of IFNγ response of cells interspersed randomly with varied densities of IFNγ-producer cells. **(b)** Comparison of population distributions of cellular IFNγ responses derived from theory and simulations. **(c)** Quantification of concordance between theory and simulations. **(d)** Histograms of MHC-I levels on IFNγ-stimulated B16 cells. **(e)** Percentages of MHC-I+ cells from (d). **(f)** Quantification of right-tailness of MHC-I levels for IFNγ-stimulated B16 cells from (g-i). Quantification of skewness of MHC-I levels from: simulations (g), B16 co-cultured with activated T cells in clusterwells (h), and B16 stimulated with IFNγ in well-mixed conditions (i).

### Supplement to figure 4

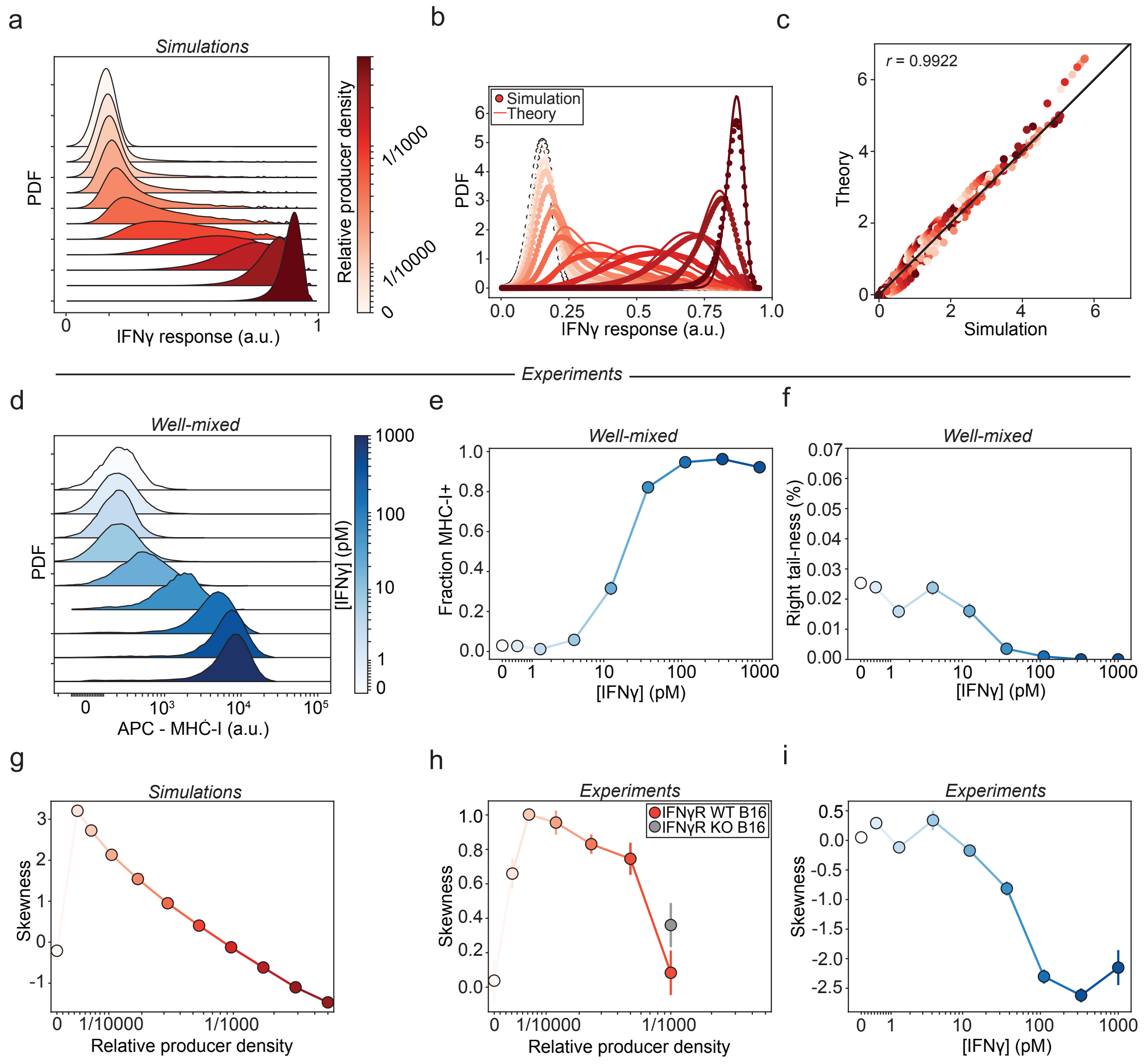

*Supplement to Figure 5: Myeloid cells are spatially correlated with regions of high TILs.*

**(a)** Image of tumor section demonstrating spatial relationship between CD11c+ myeloid cells and TILs. **(b)** Autocorrelation function of TILs, demonstrating strong, positive spatial correlation between TILs. **(c)** Cross-correlation function between TILs and CD11c+ myeloid cells, showing strong positive spatial correlation between TILs and myeloid cells. **(d-f)** Probability distribution functions for pSTAT1 **(d)**, IRF1 **(e)**, and PD-L1 **(f)** in TILs, Sox10+ melanocytes, and CD11c+ myeloid cells.

### Supplement to figure 5

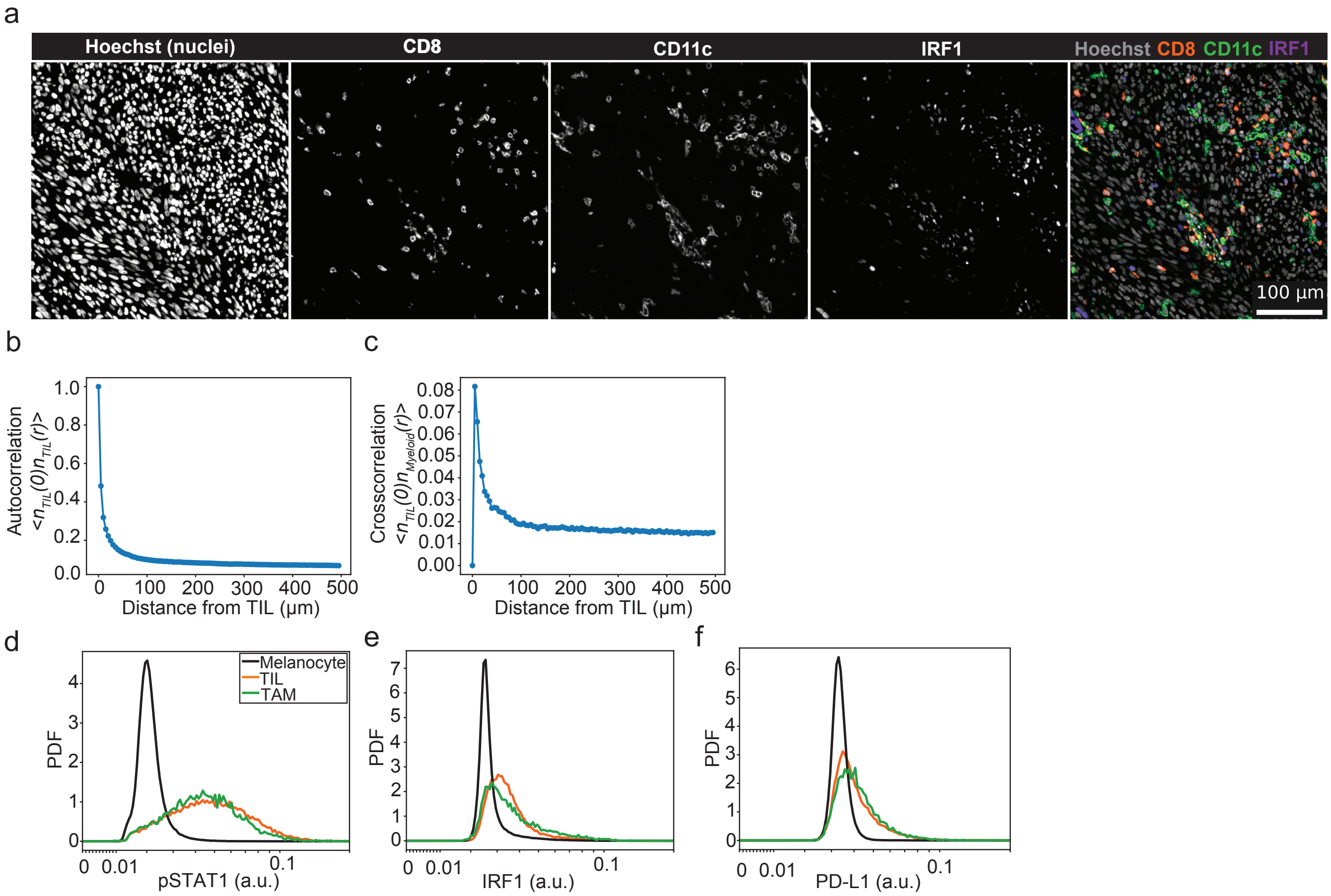

2. Mathematical framework

a. Cytokine concentration and response around single producer

As was previously described, cytokine perfusion into dense tissue can be treated as a classical diffusion-consumption problem. This can be modeled using the diffusion-consumption equation: $\frac{dc}{dt} = D\nabla^2 c - n_{consumers} k_{consumption} \frac{1}{1+\frac{1}{c}}$, where $n_{consumers}$ is the spatial density of cytokine consumers and $k_{consumption}$ is the maximal single-cell consumption rate, and $D$ the diffusion coefficient for the cytokine. The cytokine concentration $c$ is measured in units of cytokine response $EC50$. The model assumptions are only that cells do not directly affect each other's consumption ability (linearity in $n_{consumers}$).

This model brings about a length scale $\lambda_{niche} = \sqrt{\frac{D}{n_{consumers} k_{consumption}}}$ which corresponds to the typical distances cytokine will perfuse away from their source of production. Solutions to this diffusion-consumption equation show a characteristic narrow range of saturation in close proximity to the producer followed by exponential decay with length scale $\lambda_{niche}$ (Fig. 3E, S3).

Next, we will translate these single-producer response fields, into a global probability distribution of response in a population of cells.

b. Response distribution in dense tissues

We start with a dense population of cells, of which some are cytokine producers. The density of cytokine producers is $n_{producers}$ and they are assumed to be randomly distributed in space. The field around each individual producer is a spherically symmetric solution $F(r)$ of the nonlinear diffusion-consumption equation (see above). However, when these single cells are combined in a tissue, it is assumed that the cytokine fields produced by each cell can be added together by superposition. This assumption is reasonable when the density of cytokine-producing cells is low, so that the saturating regimes of cytokine concentrations do not overlap. In these cases, the concentration of cytokine produced by each individual cell can be accurately estimated and added together to calculate the total concentration in the tissue.

However, when the density of cytokine-producing cells is high, the saturating regimes of cytokine concentrations will overlap, resulting in a uniformly high response. In these cases, the exact concentration of cytokine produced by each individual cell becomes less important, as the tissue will respond with a uniform high concentration regardless. Despite this limitation, the assumption of superposition provides a useful approximation for calculating cytokine concentration in dense tissue under many conditions.

To assess the total cytokine concentration at any position in space, we need to sum over the contributions of all cytokine producers who might affect that position. We start by looking at a single volume segment $dV$ at a position $\vec{r}$ relative to our target cell. The average number of cytokine producers within this volume is $\lambda = n_{producers} dV$ and the probability of having $k$ producers within that volume follows the Poisson distribution $P_k = \frac{\lambda^k}{k!} e^{-\lambda}$. Each of these producers will contribute a concentration of $F(r)$ at the target where $r = |\vec{r}|$. The probability distribution of cytokine concentration $C$ from this volume element is therefore given by:

$$P(C, dV) = \sum_{k=0}^{\infty} \frac{(n_{producers} dV)^k}{k!} e^{-n_{producers} dV} \delta(C - kF(r)).$$

To account for the whole of space, we will need to calculate a convolution of this expression over the whole $V$. To do so, we will use the convolution theorem of Fourier transforms. We will first calculate the Fourier transform of the single volume-element expression above (this is also called the characteristic function of the distribution),

$$G(\omega, dV) = \int_0^{\infty} P(C, dV) e^{i\omega C} dC = e^{-n_{producers} dV} \sum_{k=0}^{\infty} \frac{(n_{producers} dV)^k}{k!} e^{i\omega k F(r)} = \exp(n_{producers} dV(e^{i\omega F(r)} - 1)),$$

and from that we can get the full characteristic function by integrating over space:

$$G(\omega) = \exp(n_{producers} \int_V (e^{i\omega F(r)} - 1) dV) = \exp(4\pi n_{producers} \int_0^{\infty} (e^{i\omega F(r)} - 1) r^2 dr)$$

This integral can be solved by splitting into two parts: A rapidly oscillating part in the regime $\omega F(r) \leq \pi$ where we can approximate $< e^{i\omega F(r)} >= 0$ and solve the remaining part analytically, and a smooth part where $\omega F(r) > \pi$ which can be solved numerically.

We then retrieve the full probability distribution of cytokine exposure by performing an inverse Fourier transform:

$$P(C) = \int_{-\infty}^{\infty} G(\omega) e^{-i\omega C} d\omega$$

Finally, we can translate the distribution of cytokine exposure to the distribution of cytokine response, remembering that the response to cytokine follows a hill function $R(C) = \frac{1}{1+\frac{1}{C}}$, such that $P(R) = P(C) * \frac{dC}{dR} = P(C) \cdot (C + 1)^2$.
